# *TIGRdisco*: a tool for the discovery of TIGR arrays and TIGR-associated Tas proteins from nucleotide sequences

**DOI:** 10.64898/2026.11.04.764119

**Authors:** Siwen Gu, Zheng Gong, Jenny C. Mortimer

## Abstract

**Motivation:** Tandem Interspaced Guide RNA (TIGR)–TIGR-associated protein (Tas) represents a new class of RNA-guided DNA-targeting systems, characterized by a hypercompact Tas effector guided by tigRNAs processed from TIGR arrays. In contrast to CRISPR arrays where detection relies on its near-identical direct repeats, TIGR arrays are found in dual-repeat or stem-loop architectures with extremely short box motifs and units show sequence variability within an array. This complicates its identification and limits the discovery of TIGR-Tas systems.

**Results:** Here, we present *TIGRdisco*, an integrated tool capable of discovering both the Tas protein and its associated TIGR arrays directly from unannotated nucleotide sequences. *TIGRdisco* implements independent engines to detect the two array architectures through a motif-agnostic approach. We benchmarked *TIGRdisco* against a reference dataset of TIGR-Tas loci, recovering more than 80% of array units within its search window for both types of TIGR array architectures. Furthermore, *TIGRdisco* recovered arrays for loci within clusters of Tas where associated arrays were mostly unannotated in the existing reference. We found that these arrays largely contain non-canonical box motifs or unit lengths. Lastly, as a case study, we utilized *TIGRdisco* and successfully uncovered TIGR-Tas loci across metagenomic and genomic datasets.

**Availability and Implementation:** *TIGRdisco* is open-source and available at https://github.com/JennyMortimer/TIGRdisco. All analysis outputs are attached in the Supplementary Data and available at Figshare (<u>doi.org/10.25909/34974796</u>).

## INTRODUCTION

TIGR-Tas are a recently discovered family of RNA-guided DNA-targeting systems (Faure, et al., 2025), composed of TIGR arrays and an associated, ultra-compact Tas effector protein (200–400 aa) (Liu, et al., 2026; Yang, et al., 2026). The Tas protein is structurally modular, comprising a coiled-coil (CC) domain for homodimerization and a C-terminal nucleolar protein (NOP) domain that interacts with the tigRNA and plays a role in its maturation (Zhou, et al., 2026). In addition, some Tas proteins possess N-terminal nuclease domains, categorizing them into three subtypes: Tas**R** with **R**uvC, Tas**H** with **H**NH and TasA, which lacks the nuclease domains (Faure, et al., 2025; Zhou, et al., 2026). Distinct from CRISPR (Clustered regularly interspaced short palindromic repeats) systems, the Tas effector is guided by tigRNAs processed from TIGR arrays and relies on dual spacers that pair and target both strands of DNA, independent of protospacer adjacent or target-adjacent motifs (PAMs or TAMs, respectively) (Faure, et al., 2025; Yang, et al., 2026).

Diverse Tas proteins have been computationally identified from genomic and metagenomic sources, predominantly in viruses and parasitic bacteria (Faure, et al., 2025). However, only a few have been biochemically characterized, and the TIGR arrays have not been identified in the majority of loci (Faure, et al., 2025). Being able to effectively mine and annotate complete TIGR-Tas systems in both environmental metagenomes and in genomic sequences is essential for improving our understanding of TIGR-Tas systems and enables the development of compact genome editors (Liu, et al., 2026). However, TIGR arrays are considerably more difficult to identify compared to CRISPR arrays, which consist of repetitions of a near-identical sequence gapped by unique spacers (Biswas, et al., 2016). The TIGR arrays are found in two architectures, dual-repeat (DR) or stem-loop (SL) (Figure 1A) (Faure, et al., 2025). The DR architecture is composed of a composite repetitive unit with short, alternating edge and loop repeats that flank the two spacers. These repeats show heterogeneity within the array, which complicates its detection. Meanwhile, SL architecture TIGR arrays are composed of two unique spacers linked together through a variable connector sequence and flanked by stem-forming palindromic sequences that show poor conservation within the array. Notably, both DR and SL TIGR contain conserved box C and box D motifs adjacent to spacer sequences, but these are very short (Faure, et al., 2025; Zhou, et al., 2026). Altogether, these factors present a distinctive challenge in the detection and discovery of TIGR-Tas systems where existing CRISPR array detection algorithms, developed to identify direct repeats, are likely poorly suited.

**Figure 1.**
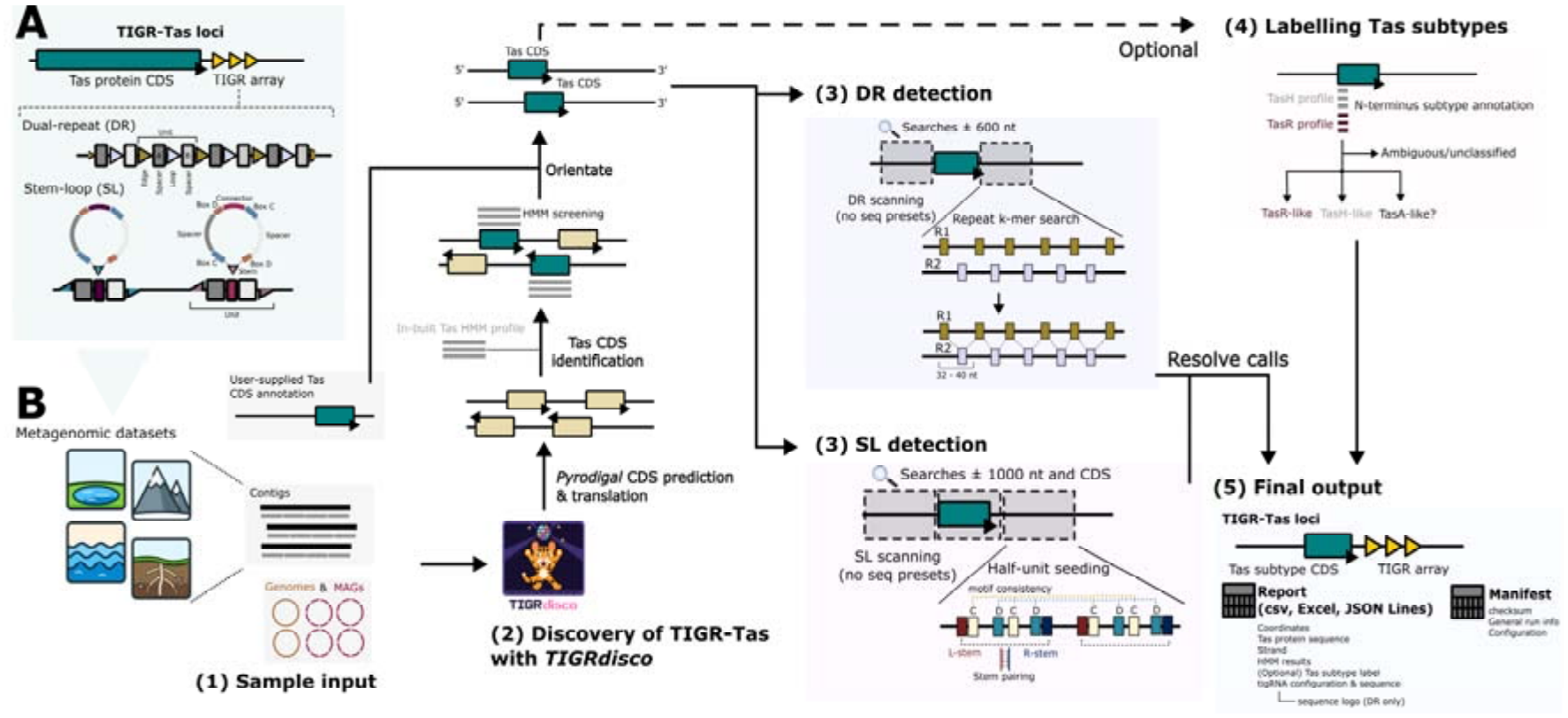
Schematic representation of TIGR-Tas loci, array architecture and the *TIGRdisco* detection workflow. (A) The TIGR-Tas locus consists of a Tas protein-coding sequence (CDS) and an adjacent TIGR array, existing in two architectures: dual-repeat (DR) and stem-loop (SL). Dual-repeat architecture consists of tandemly repeated units, each with four segments: edge repeat, spacer A, loop repeat, and spacer B. Stem-loop architecture features tandemly arranged stem loops, each with two half-units (box C–spacer–box D) joined by a connector and flanking complementary stems. (B) The five steps of *TIGRdisco*. (1) Input: metagenome, genomes or metagenome-assembled genomes (MAGs); if the user already has Tas CDS annotations, coordinates can be provided directly (db mode). (2) Anchor identification: *pyrodigal* is used to predict coding sequences, followed by screening against the built-in Tas profile HMM. (3) Independent dual-engine detection: the two engines independently scan the same locus, without any predefined motif vocabulary. (4) Tas subtype annotation (optional): N-terminal domain (NTD) of the candidate Tas is identified by hitting the NTD profile HMMs of TasH and TasR, outputting TasR-like/TasH-like/TasA-like and ambiguous/unclassified. (5) Output: the results from both engines are merged following arbitration and exported as CSV, Excel or JSON.

Therefore, we developed an all-in-one bioinformatic tool, *TIGRdisco*, for streamlined discovery of TIGR-Tas systems from user-supplied sequences and metagenomic datasets. *TIGRdisco* identifies the Tas protein coding sequence (CDS) and can also categorise detected Tas proteins into subtypes. *TIGRdisco* then employs two distinct but motif-agnostic approaches to detect adjacent TIGR arrays with DR and SL architectures. Finally, the results are reported in the output alongside the sequences and coordinates of the Tas CDS and TIGR array. We benchmarked *TIGRdisco*’s performance and applied it across multiple case studies, showing its capacity to detect and recover TIGR-Tas systems from unannotated samples.

## FEATURES AND IMPLEMENTATION

### *Overview of* TIGRdisco

*TIGRdisco* is a standalone tool for automated detection and identification of both the Tas effector and TIGR arrays from nucleotide sequences (Figure 1B). *TIGRdisco* utilizes a stepwise approach where putative Tas proteins are first identified from predicted protein coding sequences (CDS). Using the Tas CDS as an anchor, *TIGRdisco* searches for TIGR arrays in surrounding sequences and/or within the CDS. A key feature of *TIGRdisco* when detecting DR or SL arrays is that motifs are derived from the sample sequence rather than matching a predefined vocabulary, enabling more generalized detection of noncanonical TIGR-Tas systems. Furthermore, *TIGRdisco* treats each Tas CDS as an independent anchor; it can recover multiple loci per sample and detect TIGR arrays even when the search window is truncated. This makes it particularly suited for analysing metagenomic assemblies.

### Identification of Tas effector and effector subtype classification

*TIGRdisco* accepts multiple sample entry points: (1) unannotated inputs such as single sequences, contigs or metagenomic assemblies, provided in FASTA format, or (2) sequences annotated with a putative Tas CDS as a GenBank record (db mode) (Figure 1B).

For unannotated samples, *TIGRdisco* uses *Pyrodigal* to identify open reading frames (ORFs) and predict protein-coding genes (Hyatt, et al., 2010; Larralde, 2022). Using a built-in Hidden Markov Model (HMM) profile (Eddy, 2011), proteins are screened for possible Tas hits (Figure 1B). The HMM profile was constructed from known Tas proteins and covers core regions across all three Tas subtypes. Genes are considered to encode putative Tas proteins and classified into high-confidence or possible hits based on thresholds such as E-value. Regardless, each of these putative Tas CDSs is used as an independent anchor for the detection of TIGR arrays (Figure 1B).

In addition to the identification of Tas proteins, we have also built two N-termini seed profiles from known TasH and TasR classes, respectively (Figure 1B). *TIGRdisco* further analyses the N-terminus of identified Tas proteins and categorises them into either subtype (described as “TasH-like” or “TasR-like”) or otherwise grouped into “TasA-like”. Hits for Tas proteins shorter than 200 aa are not classified and assigned to an “unable to classify” category but are retained for array detection. Note that *TIGRdisco*’s subtype classification is based on HMM screening and is provisional. It also does not infer catalytic activity, and subtypes should be confirmed by structural modelling while activity needs to be biochemically tested.

### Detection of TIGR arrays with dual repeat or stem loop architectures

Before array detection, *TIGRdisco* uniformly orients the CDS of all Tas protein hits and the surrounding window (Figure 1B). Each Tas CDS is analysed by the DR and SL search engines independently. To identify DR arrays, *TIGRdisco* independently analyses a region of 600 nucleotides (nt) upstream and downstream of the anchor, allowing for truncations. The DR engine identifies periodic units consisting of two alternating repeats interspersed with spacers and filters candidates based on repeat consistency and information content, and the number and spacing of units. Within the search window, repetitive sequences are discovered using k-mer enumeration to define one repeat (R1). The second repeat (R2) is searched in a small region adjacent to the fuzzy-matched R1. This approach accommodates variation in repeat length and sequence. For called DR units, the two alternating repeats are reported as R1 and R2, which precede two spacers (referred to as A and B). *TIGRdisco* does not distinguish and assign the R1/R2 as the edge or loop repeats.

To identify SL architecture arrays, *TIGRdisco* analyses the CDS and a 1,000 nt window upstream and downstream as a whole. The CDS is included in the search, as many Tas loci were previously shown to accommodate SL units within the CDS (Xu, et al., 2026). The SL engine searches for a half-unit with a motif pair (putative box C and D) gapped by a spacer and linked to a second half-unit by a variable connector sequence. Once a full unit is identified, flanking sequences are checked for length and complementarity to form stems, considering G-U (G-T for DNA) wobble base pairing. This defines the seed SL unit. Further units are identified and added to the array based on motif matching and length consistency with the seed half-units, excluding the connector. Motif mismatches are tolerated, but recovery of those units requires longer stem complementarity.

### Data output and visualization

Upon completion of the two TIGR array detection engines, *TIGRdisco* outputs three files (Figure 1B): (1) a comma-separated values file summarizing each Tas CDS anchor, including its coordinates as well as associated array architecture, (2) a JSON Lines file documenting each Tas CDS as a record containing contig information, coordinates of CDS and array, protein and array unit sequences and HMM hit statistics, and lastly, (3) a manifest consisting of checksum, general run and configuration information. Optionally, accessible and readable *Excel* reports can also be generated for Tas loci with DR, SL or no detected arrays. The reports include contig information, the Tas and array sequences, Tas subtype classification, HMM coverage plot and sequence logos for DR arrays. In db mode with user-supplied Tas CDS annotation, coordinates and HMM hit statistics are not necessary and, hence, excluded in the outputs.

## RESULTS AND DISCUSSION

### Benchmarking

During the development of *TIGRdisco*, we noted the co-release of *TIGRFinder* and *TIGRSafari*, a tool to identify TIGR arrays and a database of annotated TIGR-Tas systems, respectively (Xu, et al., 2026). *TIGRFinder* relies on user-provided coordinates of the Tas CDS and searches for TIGR arrays. Hence, we decided to benchmark *TIGRdisco* with *TIGRFinder* on contigs provided by *TIGRSafari*, utilizing the provided Tas CDS as anchors, which included around 23,000 samples (Xu, et al., 2026).

We benchmarked the performance of the two tools on identifying TIGR arrays across the collection. *TIGRdisco* found DR arrays in 8,733 loci and SL arrays in 4,841 loci, with a total of 12,722 loci where either architecture was called. On the same input, *TIGRFinder* found 4,126 loci with DR arrays, 2,559 loci with SL arrays and 6,644 loci overall (Figure 2A). Next, using the *TIGRSafari* annotation as the reference, we mapped and tested array-level recovery. As validation of our approach, we saw 100% recovery of annotated DR and SL arrays in the fresh *TIGRFinder* run, as expected because *TIGRSafari* was built on *TIGRFinder* outputs. *TIGRdisco* conformed on 3,636 loci with DR arrays and 2,330 with SL arrays, representing 88.1% and 91.1% of *TIGRFinder* calls, respectively.

**Figure 2.**
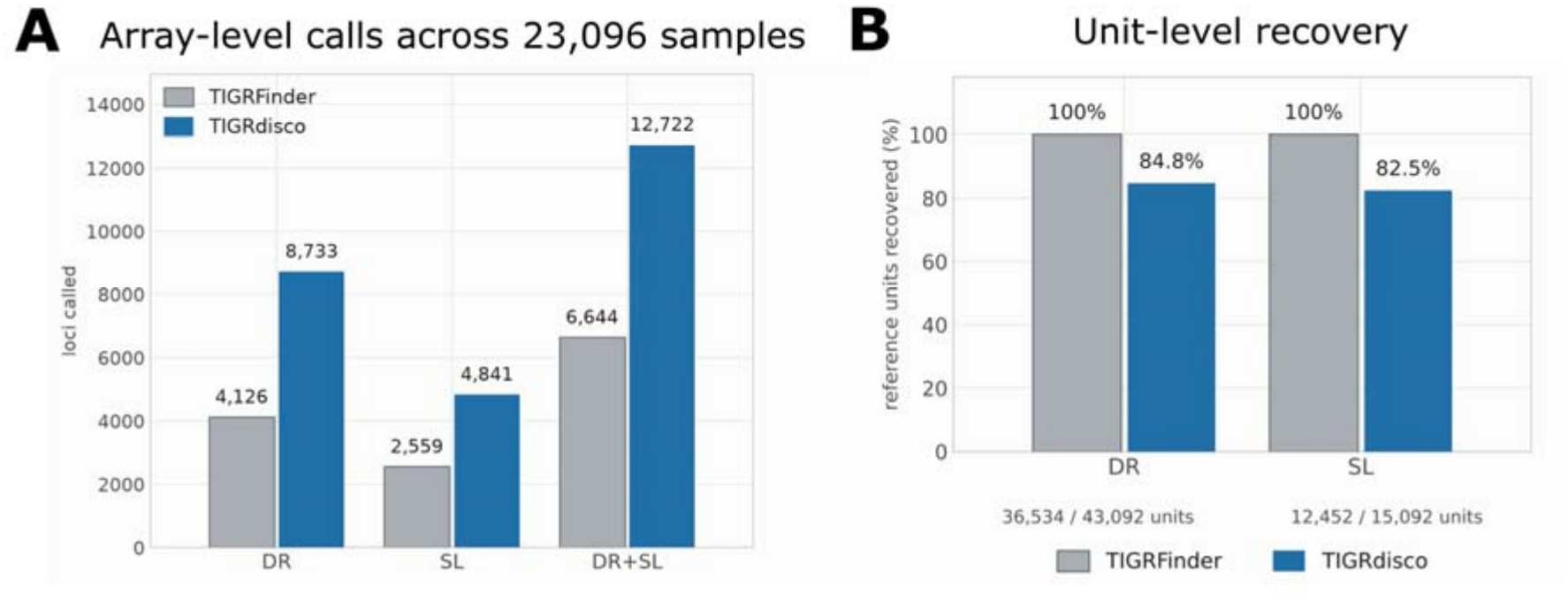
Comparison of *TIGRdisco* and *TIGRFinder* on the full reference collection. (A) Bar graph of array-level recall between the two tools of the reference from *TIGRSafari* (Xu, et al., 2026). 23,096 loci were tested and the y-axis shows the number of loci at which arrays were detected for DR, SL or the combined total (DR+SL). (B) Bar graph representing unit-level recovery between the two tools. The denominator is reference units entirely within the corresponding search window; recovery is defined as the reference unit being fully contained within a *TIGRdisco* array of the same architecture. The grey bars are the *TIGRFinder* positive control in which the reference were derived from (Xu, et al., 2026).

Next, we benchmarked *TIGRdisco* on unit-level recovery of SL and DR references. Notably, *TIGRdisco* utilizes a default 1,000 nt or 600 nt search window for SL or DR detection, respectively, and this region encompasses a majority of the annotated units (Figure S1A and S1B). Similarly, we saw 100% unit-level recovery by *TIGRFinder. TIGRdisco* recovered 82.5% of the 15,092 reference SL units within the 1,000-nt search window with 97% mapping exactly (Figure 2B). For DR array detection, *TIGRdisco* uses a 600-nt search window and recovered around 84.8% of reference units in that range (Figure 2B). The total recovery rate was around 58.7% as more than 25,000 units were outside the default detection window of *TIGRdisco*. The detection window can be extended, and we see improved recovery at the cost of runtime (data not shown).

### Uncovering the diversity of TIGR using TIGRdisco

In the *TIGRSafari* collection, Xu, et al., (2026) reported clades of clustered Tas proteins in which TIGR arrays could not be resolved using *TIGRFinder*. We analysed the length of periodic units (defined as the distance between the starting coordinates of adjacent DR units) in all DR arrays recovered by each tool, which corresponds to the length of processed tigRNA. Across approximately 8,700 Tas loci with *TIGRdisco* reported DR arrays, we found that most periodic units were the canonical 36 nt, but a greater proportion of *TIGRdisco* calls, in comparison to *TIGRFinder*, had deviating lengths of 33-35 nt and 37-39 nt (Figure 3A).

**Figure 3.**
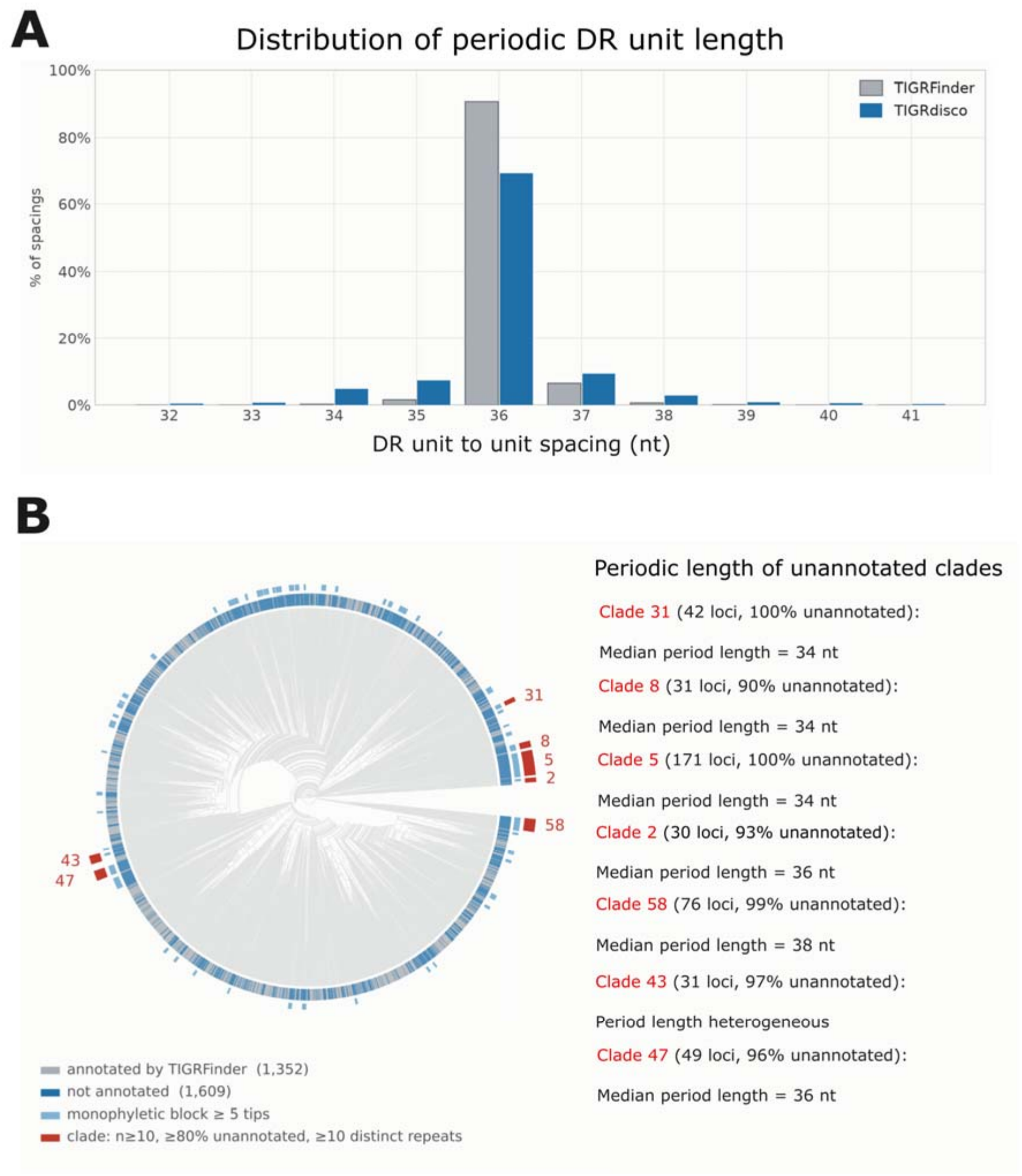
Diversity of DR TIGR arrays revealed by *TIGRdisco*. (A) Distribution of DR unit periodicity for arrays called by the two tools. The x-axis represents distance between adjacent DR unit start coordinates, whereas the y-axis represents the proportion of all inter-unit distances. The x-axis is truncated at 32-41 nt, covering 98.1% of *TIGRFinder* distances and 95.8% of *TIGRdisco* distances. (B) Under-annotated clades. The left panel shows the UPGMA dendrogram. Each tick on the outer ring represents one non-redundant representative protein. Grey represents *TIGRFinder* array annotation, blue represents no annotation. Light blue highlights monophyletic blocks of ≥ 5 tips. Red highlights the seven under-annotated branches (under a K = 64 cut, branches with ≥ 10 representatives of which ≥ 80% are unannotated). The right panel shows the number of loci, the unannotated proportion of loci and the median inter-unit spacing for the seven branches.

To further investigate unresolved loci, we constructed a dendrogram from nearly 3,000 representative Tas proteins. These non-redundant representatives were derived from 8,686 loci with *TIGRdisco-*annotated DR arrays, excluding ones with short Tas proteins. The dendrogram was cut into 64 deep branches to identify 7 branches that are substantially under-annotated by *TIGRFinder* (Figure 3B). We then expanded representatives to their cluster members and analysed the 430 loci of these branches. The average intra-branch protein identity was 47.6% compared to 26.2% between branches, and cutting into 32 or 128 deep branches resulted in 6 or 8 of these branches, respectively. Re-running *TIGRFinder* across the 7 branches only recovered arrays in around 2.1% of the members.

*TIGRdisco*’s motif-agnostic and permissive approach reveals two independent potential causes. First, 4 of the 7 branches had median periodic unit lengths of 34 or 38 nt, and 1 branch was heterogeneous. Second, we observed many arrays with units consisting of non-canonical box C/D motifs outside the *TIGRFinder* expected sequence. Units of some arrays deviate from both the canonical unit length and motif. These unconventional arrays should be further experimentally tested to confirm our findings. We also found arrays that conform to *TIGRFinder*’s length and motif restrictions but were still unannotated, implying other underlying reasons that call for further investigation. The *TIGRdisco* annotation of these loci is provided in the Supplementary Data.

*TIGRFinder* uses sequence matching with a preset box C/D motif vocabulary and length constraints to identify units of motifs flanking a spacer (Xu, et al., 2026). This strategy allows high-confidence recovery of units that fit the vocabulary but may miss ones that deviate, potentially representing divergent systems. Even though *TIGRdisco* identified variable periodic unit length, it cannot recover any arrays outside its constraints on inter-unit spacing (32–40 nt). We also expect reduced confidence in recovery accuracy compared to *TIGRFinder* as compensation for increased tolerance to sequence variation, particularly for SL arrays that lack definitive repetitive sequences. Despite these constraints, *TIGRdisco* uncovered loci of TIGR-Tas systems with unconventional arrays, showing unexplored diversity beyond known features.

### Discovery of TIGR-Tas systems from metagenome datasets

To demonstrate the utility of *TIGRdisco* in analysing metagenomes, we deployed it to search for novel TIGR-Tas systems in datasets derived from underwater hydrothermal sediment and from a hot spring in Yellowstone National Park. *TIGRdisco* was capable of processing the datasets and identified multiple TIGR-Tas systems. We refer to these TIGR systems as HydroTas and SpringTas, which correspond to the environments. To confirm these results, we first selected a TIGR-Tas system (HydroTasA-1) that was labelled as “TasA-like” by *TIGRdisco*. Using *ESMFold2* (Candido, et al., 2026) to predict its structure, we observed the signature CC linked to a C-terminal NOP fold (Figure 4). This is further supported by the extracted Sparse autoencoder (SAE) features using *ESMC* (Candido, et al., 2026), where “NOP56/58-PRP31 central coiled-coil” and “C-terminal nucleic acid-binding module” were activated (Figure 4). Analysis of two other DR-associated (HydroTasA-2 and SpringTasA-1; Supplementary Data) and one SL-associated (SpringTasA-2; Figure 4) “TasA-like” candidates showed the expected structural folds and SAE feature activation.

**Figure 4.**
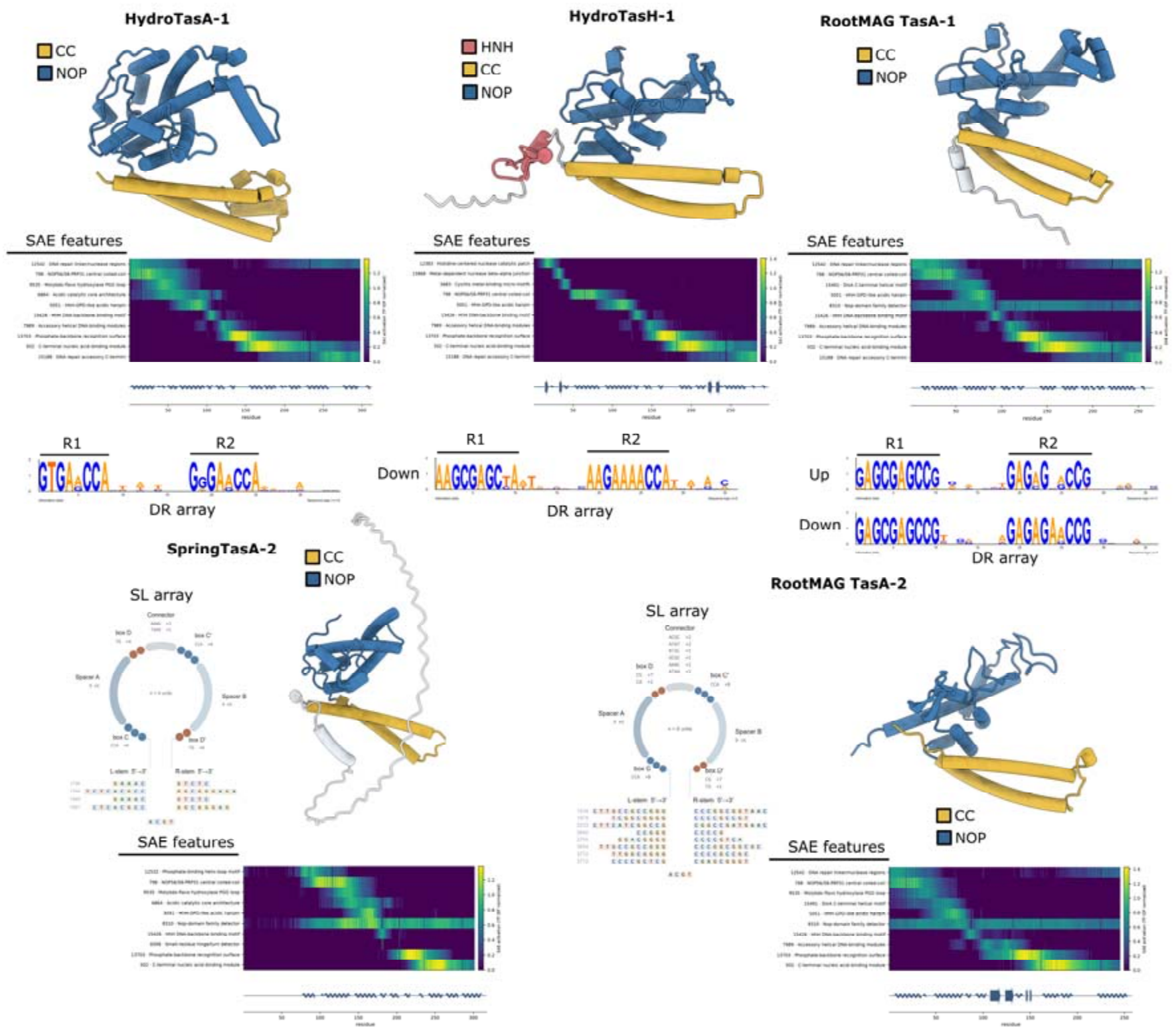
Mining TIGR-Tas systems across case studies. Five representative TIGR-Tas systems from hydrothermal vent (Hydro), Yellowstone National Park hotspring (Spring) metagenomes, and root rhizosphere MAGs (RootMAG) are shown. The protein structure was predicted using *ESMFold2* and rendered in *ChimeraX*. CC, coiled-coil. Heatmap showing the activation of top 10 sparse autoencoder (SAE) features for each Tas sequence extracted using *ESMC* and a linear positional schematic of the predicted protein secondary structure constructed using *mkdssp* assignment. For DR arrays (HydroTasA-1, HydroTasH-1 and RootMAG TasA-1), the sequence logo from *TIGRdisco* output is directly displayed with R1 and R2 labelled. Logo for arrays upstream (‘Up’) and downstream (‘Down’) of the Tas CDS is shown when relevant. For HydroTasH-1, only the downstream arrays are shown as the upstream array did not meet the *TIGRdisco* threshold for logo construction. For SL arrays (SpringTasA-2 and RootMAG TasA-2), a representative stem loop structure with motif, connector and stem sequences labelled. All *TIGRdisco* outputs for these representatives and other identified TIGR-Tas systems are included in the Supplementary.

Next, we focused on a high-confidence Tas candidate (HydroTasH-1) labelled as “TasH-like”. Structural prediction showed the HNH nuclease, CC and NOP folds (Figure 4). Extracted SAE features show additional activation of the “Histidine-centered nuclease catalytic patch”. HydroTasH-1 was associated with upstream and downstream DR TIGR arrays. Interestingly, we noticed that the HydroTasH-1 CDS was annotated in the *TIGRSafari* database but without TIGR arrays (Xu, et al., 2026). We used *TIGRFinder* to search the contig and were also unable to detect any arrays. This may be because one of the two repeats varied between a CCA and CTA box C motif, showing the within-array heterogeneity of TIGR systems. Overall, these results demonstrate the application of *TIGRdisco* in mining TIGR-Tas systems from metagenomes. The *TIGRdisco* outputs and curated amino acid sequences of analysed Tas effectors are included in the Supplementary Data.

### Discovery of TIGR-Tas systems from genomes and metagenomic-assembled genomes (MAGs)

Identifying TIGR-Tas systems from assembled genomes is fundamental to understand their evolutionary relationship and function. We further used *TIGRdisco* for the identification of TIGR-Tas systems in genomes of phage, archaea as well as in metagenome-assembled genomes (MAGs). We first employed *TIGRdisco* to search the *Salicola* phage CGphi29 genome from which TIGR-SpTasH was derived (Faure, et al., 2025). *TIGRdisco* specifically identified SpTasH and its associated TIGR array.

Next, we investigated the capacity of *TIGRdisco* to process and search datasets of MAGs. We deployed *TIGRdisco* to analyse a collection of MAGs derived from plant root rhizosphere. We identified 14 putative Tas proteins labelled as high confidence with associated TIGR arrays, 5 of which with DR and 9 with SL architectures. We selected two TasA effectors (RootMAG TasA-1 and RootMAG TasA-2) for structural prediction as described previously. In both, we saw similar TasA structural domains and activation of those SAE features (Figure 4). Overall, this case study showed the capacity of *TIGRdisco* in detecting TIGR-Tas loci from genomic datasets.

## LIMITATIONS AND CONCLUSION

There are several limitations of *TIGRdisco*. Because *TIGRdisco* does not impose predefined box motif vocabulary, the higher unit recovery yield may include false positives, particularly in scenarios with extreme GC content. Second, *TIGRdisco* uses a default search range from the Tas CDS anchor and misses ones outside. In addition, the DR array search engine is unable to classify edge and loop repeats as these distinctions are dependent on which repeat is processed while the other is internal during tigRNA maturation (Faure, et al., 2025; Xu, et al., 2026). *TIGRFinder* utilizes positional nucleotide complementarity as a parameter to classify the edge repeat (Xu, et al., 2026). Therefore, we recommend harnessing both approaches and their tools for improved discovery of TIGR-Tas systems from sequence data.

In conclusion, to overcome the challenges and limited available tools to mine TIGR-Tas systems, we developed *TIGRdisco*, a command-line tool which combines both Tas protein identification and TIGR array detection. *TIGRdisco* is open-source and accessible through GitHub (available upon publication). We benchmarked *TIGRdisco* and tested it across metagenomic and genomic datasets. *TIGRdisco* recovered arrays across Tas loci that were previously unannotated, revealing the diversity of TIGR arrays with canonical and non-canonical architectures.

## METHODS

### Construction of Tas protein HMM profiles

HMM profiles of Tas proteins were constructed using *PyHMMER* and *hmmbuild* on Galaxy (Eddy, 2011; Eddy, 2023; Larralde and Zeller, 2023; The Galaxy, 2026). Three HMM profiles were used in *TIGRdisco*: (1) the core Tas HMM profile used to screen for candidate proteins, (2) the N-terminal domain (NTD) profile HMM of TasR, and (3) TasH. We used Tas proteins derived from Faure, et al., (2025) for initial clustering using *MMseqs2* (Steinegger and Söding, 2017) with 80% sequence identity and 80% target sequence coverage (cov-mode 1) and filtered for sequences with length ≥ 200 aa. The remaining 5,253 sequences were aligned with *MAFFT* (-- auto mode) (Katoh, et al., 2002) and used to construct the HMM profile using *hmmbuild* with default parameters. This resulted in the core Tas HMM profile, MITTas.hmm3 (357 match states). Based on the HNH and RuvC labels in NTD_REP50 of Faure, et al., (2025), TasH, TasR and TasA were extracted. Following sequence clustering and processing as described, 189 TasH, 774 TasR and 4,290 TasA sequences were retained from each subtype, respectively. After performing alignment and HMM construction on TasA sequences, we obtained MITTasA.hmm3 (337 match states). This profile was used to identify the conserved Tas core and hit the best-scoring domain for TasH and TasR (independent domain E-value ≤ 10^-5^, HMM coverage ≥ 50%). The N-terminal sequences from 30 to 200 aa before the alignment start position were extracted, and redundancy was reduced again with *MMseqs2*. The remaining N-terminal sequences were then subjected to multiple sequence alignment using *MAFFT* L-INS-i, then applied to construct models using *PyHMMER* with the ‘fast’ mode and default parameters. This resulted in TasH_NTD_seed.hmm (107 match states) and TasR_NTD_seed.hmm (154 match states).

### Annotation of input samples, identification and subtype classification of Tas CDS

For unannotated samples, *TIGRdisco* requires a FASTA nucleotide input. ORFs are identified using *Pyrodigal* in metagenomic mode. Translated proteins of at least 60 aa and without internal stop codons are retained for analysis. The ORFs are screened for Tas candidates using *PyHMMER* with the built-in core Tas HMM profile. Hits with a full-sequence E-value of 1x10^-5^, score of at least 5 bits, HMM coverage of at least 20%, protein coverage of at least 8% and an independent E-value of less than 1 are considered passing candidates. The CDS of these proteins are retained as independent anchors for TIGR array detection. Passing candidate proteins are labelled as “high confidence” with full-sequence E-value lower than 1x10^-15^, score of at least 50 bits, HMM coverage greater than 40%, protein coverage greater than 45% and independent E-value lower than 1x10^-8^ or are otherwise labelled as a possible hit category. To annotate the Tas subtypes of hits, the TasH and TasR N-terminus HMM profiles are used to screen candidate proteins that are at least 200 aa. To be classified as “TasH-like”, a domain score greater than 48 bits or an independent E-value lower than 1x10^-5^ is required; “TasR-like” categorisation requires a score greater than 58 bits or an independent E-value lower than 1x10^-5^. Dual-positive proteins are labelled as “ambiguous”, and proteins passing neither profile are labelled “TasA-like”. “TasA-like” is a profile-based label, not proof that a nuclease domain is absent. For annotated samples, a GenBank record with a “CENTRAL” label representing the putative Tas protein of interest is supplied, bypassing ORF prediction and HMM screening.

### TIGR array detection

All Tas CDS are treated as independent anchors for TIGR array detection. All anchors are oriented, and search windows are clipped at record boundaries. The search window of each anchor is analysed with the DR and SL detection engines separately.

The DR detection engine searches 600 nt flanking the Tas CDS. The upstream and downstream searches are independent. Within each 600-nt flanking region, all k-mers of lengths 8-13 nt are enumerated and sorted by the occurrence count and length of products. The top 15 are selected as candidates for R1 and paired with R2, an equal-length substring 7–13 nt downstream of the matching site, for array construction. For each candidate pair (R1, R2; count ≥ 8), the first R1 is located via approximate matching and serves as an anchor. Then the process proceeds unit by unit following the structure, R1 – spacer A – R2 – spacer B, until no further R1 can be detected. The units obtained from the above progression are defined as “Core”. As the progression begins at the anchor, there may still be uncovered divergent units at both ends of the array. Therefore, an iterative scan is performed in the upstream (“Prefix”) and downstream (“Suffix”): whenever an uncovered R1 is detected, it is used as a new front to continue the search outwards until no matches are found. The array is scored based on the sum of the information content per column of the aligned repeats, and the quantity is displayed in the sequence logo. Scoring of column:

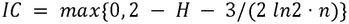

H□ is the Shannon entropy of the i-th column, n is the number of sequences, and the final term represents a small-sample correction. The score for an array is the sum of the column-wise information content for R1 and R2 in the aligned sequences. This score is used both to determine the unit register with the highest information content and for subsequent filtering. A candidate array must contain more than three valid units and at least 18 bits of information content. It must also satisfy periodicity requirements: in a sliding window of four consecutive units, the median of the three inter-unit spacings must fall within 32–40 nt, with at least 75% of spacings differing by no more than 3 nt. Among the suitable candidates, the pair (R1, R2) with the highest information content is retained as the output array for that flank. The SL detection engine searches within the CDS and flanking 1,000 nt. An SL unit is defined as a seven-field structure of box C – spacer A – box D – connector – box C – spacer B – box D, enclosed by a stem. The SL engine enumerates all the box C and box D (2–4 nt) and an equal-length spacer (8–10 nt) between them, then searches for an exact repeat structure of box C – spacer – box D at a distance of 2–20 nt (connector) within the window. The stem extends outwards from both ends of the unit via Watson–Crick base pairing, accommodating G-T pairing (G-U wobble in RNA), and permits at most one mismatch. The stem length at the seeding site must be ≥ 7 nt. The SL engine then searches for additional units matching the same seed motif (C, D) within a radius of 1,500 nt. At this stage, spacers A and B are independently enumerated as 8–10 nt without an equal-length requirement. Motif (C, D) match is balanced between the required stem length and error tolerance. The requirement for stem length gradually becomes more stringent with the increase of mismatches across the four copies. The total frame length of 2(|box C| + |box D|) + |spacer A| + |spacer B| within the same array remains consistent. Units are clustered based on a spacing of ≤ 300 nt. The clusters are defined as a single array with at least 3 units. The detection process does not rely on any predefined motifs. Both box C and box D are discovered as spontaneously repeating motifs within the window. On this basis, a re-scan of positive sites is performed. The first scan enumerates length-free boxes and determines the positive sites with arrays. For sites judged positive in the first scan, the same window is re-read with a frame of |box C| = 3 and |box D| = 2. If this frame also yields a valid array, that reading is adopted. Otherwise, the result from the first scan is retained. Once both engines are completed, an arbitration is applied for each anchor. An SL array is removed only when all units overlap at least one DR unit by one or more base pair. The DR calls remain unchanged.

### Benchmarking

To benchmark *TIGRdisco* with *TIGRFinder*, the reference annotation of TIGR-Tas loci was sourced from *TIGRSafari* (Xu, et al., 2026). *TIGRFinder* (version 1.0.0) was used with default parameters for a fresh run, which reproduced the reference annotations exactly, with no discordance at any locus. The same records were then processed with *TIGRdisco* (version 1.2.4) in db mode, which annotates arrays from GenBank with “CENTRAL” feature as the Tas CDS anchor. Coordinates from both sources were converted to 0-based half-open intervals on the original records. The reference annotation’s GenBank feature locations, which are 1-based inclusive intervals, were converted to [start - 1, end). Annotation records were reverse-complemented when the “CENTRAL” feature carried a complement location, and the coordinates were subsequently mapped back to the original records. Reference unit intervals were retrieved from the annotated GenBank features. Intervals of DR units detected by *TIGRdisco* were defined by the starting coordinates and summed length of their sequence components (R1 + spacer A + R2 + spacer B). SL unit intervals were defined by the start and end coordinates. Instead of pairwise comparison between unit intervals from *TIGRdisco* and the reference, we scored recovery against the union of all *TIGRdisco* unit intervals. This is because the two tools define boundaries at offset positions for DR array units. For consistency, this was applied when benchmarking both the DR and SL array detection engines. Reference units are recovered when all of their nucleotides are captured by the union. Recovery was reported as two rates: (1) within-window rate, defined as the reference units recovered by *TIGRdisco* divided by all reference units within the search window, and (2) overall recovery rate, which includes reference units outside the window. The search window was defined by the coordinates of the “CENTRAL” feature.

### Periodic unit length variation

At reference loci with DR arrays that *TIGRdisco* identified, the periodic unit length was defined as the median length of complete units in the array. The length was determined as the distance between starting coordinates of adjacent DR units. To eliminate pseudoreplication, redundancy was removed based on the anchored protein sequences. The set of amino-acid 5-mers was constructed, and the standard Jaccard coefficient was used as a similarity measure; proteins were greedily clustered at a threshold of 0.45. The first retained entry in each cluster was regarded as the representative, yielding 2,961 non-redundant representatives. Periodic unit lengths were calculated for both the loci of non-redundant and unclustered sets, confirming that distribution was not affected by clustering.

### Analysis of unannotated clades

The 2,961 non-redundant representative sequences obtained from previous processing were categorised based on their *TIGRFinder* array annotation (Xu, et al., 2026). 54.3% (1,609) non-redundant representative sequences were unannotated. All non-redundant representative sequences were aligned against the MITTas.hmm3 using *hmmalign* (Eddy, 2023; Larralde and Zeller, 2023), extracting the 357 match-state columns. Identity was calculated for ungapped columns, with distance defined as 1 − identity. A dendrogram was constructed with the Unweighted Pair Group Method with Arithmetic Mean (UPGMA). The dendrogram was cut at K = 64 (fcluster, maxclust), yielding 16 branches with ≥10 sequences. Seven branches, each with ≥10 representative sequences, were under-annotated, defined as ≥80% of sequences within the branch unannotated. Each representative sequence was also expanded to its protein cluster, and *TIGRFinder* was re-run with the default settings to detect arrays. Cuts of K = 32 and K = 128 were also performed. *TIGRdisco* outputs of loci in these branches were extracted from the previous benchmarking run. These were used to curate the Supplementary Data which was examined manually to identify possible causes of under-annotation. The periodic unit length, as defined above, was used to first calculate within array median and was then used to determine the median length across arrays in the branch. Fitch parsimony was performed on array annotation status of the dendrogram and required at least 560 state changes. After 1,000 tip-label permutations, the null distribution was 975.1 ± 15.3 (minimum 915).

### Mining TIGR-Tas systems in case studies

The Dombrowski hydrothermal vent sediment metagenomes, *Salicola* phage CGphi29 genome (NC_020844.1), the JGI microbial mat metagenomes (Yellowstone National Park), and the Xu citrus rhizosphere MAG were analysed using *TIGRdisco* in FASTA mode with the default detection parameters. The Dombrowski, JGI, and Xu datasets were divided into 53, 212, and 223 input batches, respectively, without truncating sequences. Each batch was searched independently, and the results were subsequently combined. Therefore, HMM E-values reflect the search space of each individual batch. The complete phage genome was analysed as a single input. No limit on the number of input records was applied in any analysis.

### Structural prediction and SAE feature extraction

All structural predictions were conducted using *ESMFold2* (Candido, et al., 2026) with the model esmfold2-fast-2026-05 and default settings. The protein sequence was analysed using *ESMC* (Candido, et al., 2026) (esmc-6b-2024-12 model) to extract SAE features. Maximum activation across all positions was used to rank features, and the top 10 highest SAE features and their labels were extracted.

## DATA AND CODE AVAILABILITY

*TIGRdisco* is open-source and freely available through Github (https://github.com/JennyMortimer/TIGRdisco). Analysis of *TIGRdisco* outputs at branches consisting of unannotated TIGR-Tas loci were deposited in Figshare (doi.org/10.25909/34974796). *TIGRdisco* outputs of case studies were also deposited in Figshare. Additional information available in the Supplementary Data. Sequence and *TIGRdisco* output of HydroTas, SpringTas and RootMAG Tas is available in the Supplementary Data. Contigs with annotated TIGR-Tas systems were retrieved from *TIGRSafari* (Xu, et al., 2026). The metagenome dataset from hydrothermal vent was retrieved from the SPIRE database (https://spire.embl.de/spire/v1/study/Dombrowski_2018_sediment). The metagenome dataset from Yellowstone National Park was retrieved from SPIRE database (https://spire.embl.de/spire/v1/study/JGI_microbialmat_USA_Mexico). The root rhizosphere MAGs were retrieved from the SPIRE database (https://spire.embl.de/spire/v1/study/Xu_2018_citrus). The *Salicola* phage CGphi29 genome was retrieved from NCBI (accession: NC_020844.1).

## AUTHOR CONTRIBUTIONS

Siwen Gu (Conceptualization [equal], Data curation [lead], Formal analysis [lead], Methodology [equal], Software [lead], Writing – original draft [supporting], Writing – review & editing [equal]), Zheng Gong (Conceptualization [equal], Formal analysis [supporting], Methodology [equal], Supervision [equal], Writing – original draft [lead], Writing – review & editing [equal]), Jenny C. Mortimer (Conceptualization [equal], Methodology [equal], Supervision [equal], Funding acquisition [lead], Writing – review & editing [equal]).

## ACKNOWLEDGMENTS

We acknowledge Prof. Matthew Gilliham for his suggestions on this work. We also thank all members of the Mortimer lab and ARC CoE in Plants For Space for their support. Generative AI tools (Anthropic’s Claude Code, Opus 5) were used to generate and refine portions of the code based on the authors’ design and input. All code was reviewed, tested and validated by the authors. The authors take full responsibility for the software and the results reported in this manuscript.

## FUNDING

This work was funded by the Australian Research Council (ARC) as part of the Centre of Excellence in Plants For Space (P4S; CE230100015).

## Notes

https://doi.org/10.25909/34974796

### Competing Interest Statement

The authors have declared no competing interest.

### Summary of Updates

Revised abstract and main text. Supplemental files updated.

