## Supplementary Information for "*TIGRdisco*: a tool for the discovery of TIGR arrays and TIGR-associated Tas proteins from nucleotide sequences"

**SUPPLEMENTARY FIGURES**

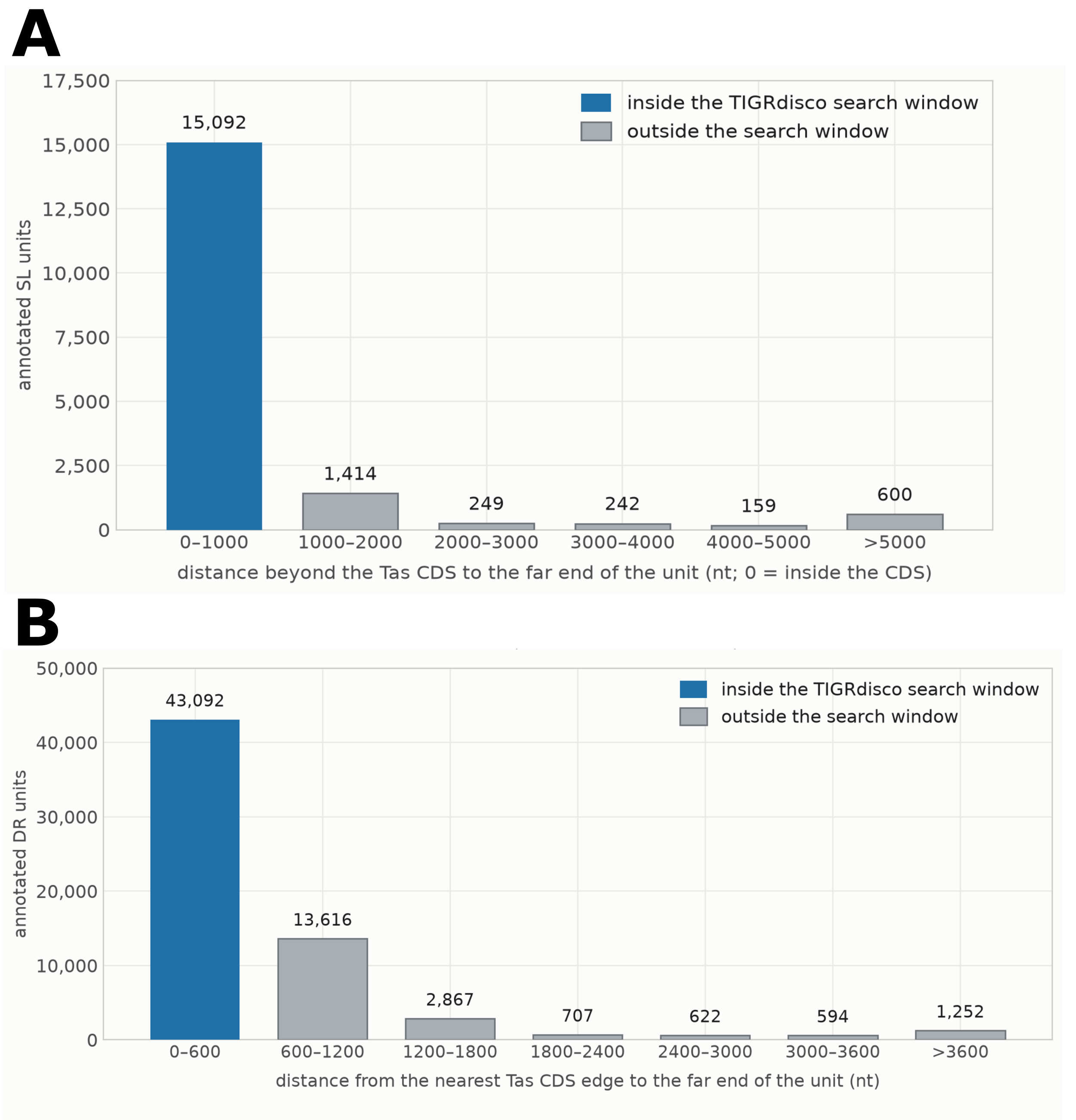

**Figure S1.** Analysis of TIGR array unit distances from the Tas CDS anchor.

(A) Distribution of distances between annotated SL units and the Tas CDS. All *TIGRSafari* reference SL units were binned by distance beyond the Tas CDS. Those within the CDS were assigned 0. The *TIGRdisco* SL window spans the Tas CDS ±1,000 nt and includes the CDS itself. The blue bar represents the units falling inside the *TIGRdisco* search window. The grey bar denotes units outside the window. The bin on the far right is opened.
(B) Distribution of distances between annotated DR units and Tas CDS. All *TIGRSafari* reference DR units are binned by distance beyond the Tas CDS. The *TIGRdisco* DR window spans two 600-nt flanking regions and excludes the CDS itself. The blue bar represents the units falling inside the *TIGRdisco* search window. The grey bar denotes units outside the window. The bin on the far right is opened. The bin on the far right is opened. A further 2,885 units overlap the Tas CDS and were not plotted.
